# Temperature-dependent reprogramming of virulence traits during *Pseudomonas aeruginosa* biofilm formation

**DOI:** 10.64898/2026.09.16.751985

**Authors:** Karishma Bisht, Alex R. Luecke, Catherine A. Wakeman

## Abstract

*Pseudomonas aeruginosa* is an opportunistic pathogen that occupies diverse ecological niches, including soil, water, and the human host. Environmental cues encountered across these habitats trigger adaptive responses that promote survival and persistence through regulation of virulence-associated traits, including secreted factors, siderophores, and biofilm exopolysaccharides (EPS). One major change experienced during the transition from environmental reservoirs to the host is an increase in temperature. Although temperature is a key signal encountered during host transition, its global impact on *P. aeruginosa* physiology remains poorly understood. We therefore investigated the effects of temperature on planktonic and biofilm-populations, comparing both growth states at 23°C and 30°C, representing environmental temperatures, and at 37°C and 40°C, representing normal and febrile host temperatures. Transcriptomic and phenotypic analyses revealed extensive temperature-dependent regulation of virulence determinants in both growth states. Expression of the type VI secretion system was elevated at environmental temperatures, whereas pyoverdine biosynthesis and the type III secretion system were upregulated at host temperatures. Building on our previous finding that biofilms formed at environmental and host temperatures differ in architecture, biomass, and EPS composition, we next examined how these structural differences influence stress tolerance. Biofilms grown at 23°C and 30°C exhibited substantially greater tolerance to antibiotic stress than biofilms grown at 37°C and 40°C. Growth temperature therefore establishes biofilm properties that subsequently influence the stress-tolerance profile of the population. Collectively, our findings identify temperature as a major environmental cue that reprograms *P. aeruginosa* physiology in ways likely to support persistence across distinct ecological niches.

**Importance:** *Pseudomonas aeruginosa* causes serious infections in individuals with cystic fibrosis, burn wounds, and compromised immune systems, but also persists in diverse environmental reservoirs. During transition from the environment to the host, this bacterium encounters a predictable increase in temperature that may regulate physiology and virulence. Because most laboratory studies are conducted at 37°C, the effects of environmentally relevant temperatures remain poorly understood. We examined gene expression and virulence-associated phenotypes across four temperatures in both planktonic and biofilm growth states. Temperature differentially regulated host-associated and environmental virulence programs, including elevated T3SS at host temperatures and T6SS at environmental temperatures. Notably, biofilms formed at environmental temperatures exhibited greater antibiotic tolerance. These findings reveal temperature as a powerful ecological cue directing environmental-to-host adaptation in *P. aeruginosa*.

## Introduction

*Pseudomonas aeruginosa* is a ubiquitous gram-negative proteobacterium capable of colonizing diverse niches, ranging from fresh water and soil environments to the immunocompromised human host (1). It also persists on and within hospital surfaces, such as contaminated sinks, making it a pervasive source of nosocomial infection (2, 3). This opportunistic pathogen establishes successful infection and colonization across a wide range of environment in part by secreting several virulence factors (4–7). Additionally, its ability to form biofilms makes it more tolerant to both the antibiotic treatment and the host immune response than its planktonic counterpart (8, 9). Together, these traits can not only result in severe wound infections but can also increase both the morbidity and mortality rate in several chronic infection associated diseases, such as cystic fibrosis (CF) and chronic obstructive pulmonary disease (COPD) (10, 11).

Temperature is a key environmental signal for bacteria that transition between external reservoirs and mammalian hosts, and many pathogens couple thermal shift to the expression of virulence determinants (12). In *P. aeruginosa*, this sensing occurs through several mechanisms, including RNA thermometers, in which structured 5′ untranslated regions melt at elevated temperature to expose ribosome binding sites (13, 14); temperature-sensitive protein activity (15); and thermally responsive second messenger signaling (16). Consistent with this, transcriptomic comparisons of ambient and host temperature indicate that a substantial fraction of the genome is thermoregulated (17–19). Importantly, this response is not fixed but depends on physiological state, as temperature has been shown to regulate distinct sets of genes at different growth phases (19, 20). Which genes respond to a thermal shift therefore depends on the state of the cell, yet thermoregulation has been examined almost exclusively in planktonic culture and across a narrow temperature range, leaving its behavior in the biofilm lifestyle and across the wider thermal range *P. aeruginosa* encounters largely undefined.

We have previously shown that thermoregulation reshapes the genetic circuitry of *P. aeruginosa* affecting its biofilm properties, in part through temperature-dependent proteome remodeling that elevates the expression of Pf1 phage proteins. This protein promotes more robust biofilm formation at 37°C, thereby helping *P. aeruginosa* adapt to the mammalian host (21). Given that Pf1 phage proteins may themselves act as virulence determinants, we reasoned that temperature may coordinate a broader virulence program.

To test this, we used RNA sequencing to compare the transcriptome of *P. aeruginosa* strain UCBPP-PA14 at 23°C and 30°C, mimicking environmental and industrial temperatures, and at 37°C and 40°C, mimicking normal and febrile host body temperature, in both biofilm and planktonic growth states. Our results indicate that temperature differentially regulates virulence factor production in both growth states, facilitating *P. aeruginosa* adaptation to and transition into the human host. Differentially expressed genes included those encoding the type III and type VI secretion systems, as well as quorum sensing-regulated products such as pyocyanin and the siderophore pyoverdine. To assess the functional implications of these transcriptional changes, we examined biofilm formation across the four temperatures by measuring biofilm biomass. We found that while some genes are essential at all the temperatures, others are more temperature restricted. Consistent with temperature-dependent differences in biofilm properties previously reported by our group and others, as well as with observed differences in biofilm colony morphology, we found corresponding differences in tolerance to stressors commonly used to target biofilms. Although *P. aeruginosa* transcriptomic response to temperature have been reported previously (17, 18, 22–25), to our knowledge this is the first study to span a range of temperatures representing both environmental and host conditions in both planktonic and biofilm growth states.

## Results

### Growth state and temperature distinctly alter *Pseudomonas aeruginosa* transcriptome

Our previous work showed that the biofilm proteome of *P. aeruginosa* is highly impacted by temperature shifts (21). We therefore wanted to determine some of the specific gene expression changes that might contribute to temperature-driven virulence factor production. We performed transcriptomic analyses on both planktonic and biofilm-associated cells at 23°C and 30°C mimicking environmental/industrial temperatures and 37°C and 40°C mimicking host body temperature under normal and febrile conditions **(Fig. 1A).** Principal component analysis for *P. aeruginosa* PA14 strain grown in biofilm and planktonic form at all the four temperatures revealed a notable clustering effect of the replicates for biofilm as well as the planktonic stage **(Fig. 1B)**. It showed that the transcriptomes tend to cluster near one another based on the growth state, with the biofilm transcriptome at 40°C most distinct from the other samples. Genes were differentially expressed in both the planktonic and biofilm state at all the four temperatures. The selection criteria for differential expression required genes to have a fold change of ≥2 and a q value of ≤0.05 to be considered significant. We were also able to categorize the number of unique and shared, differentially expressed genes across temperature conditions in biofilm versus planktonic growth state **(Fig. S1)**

**Fig. 1.**
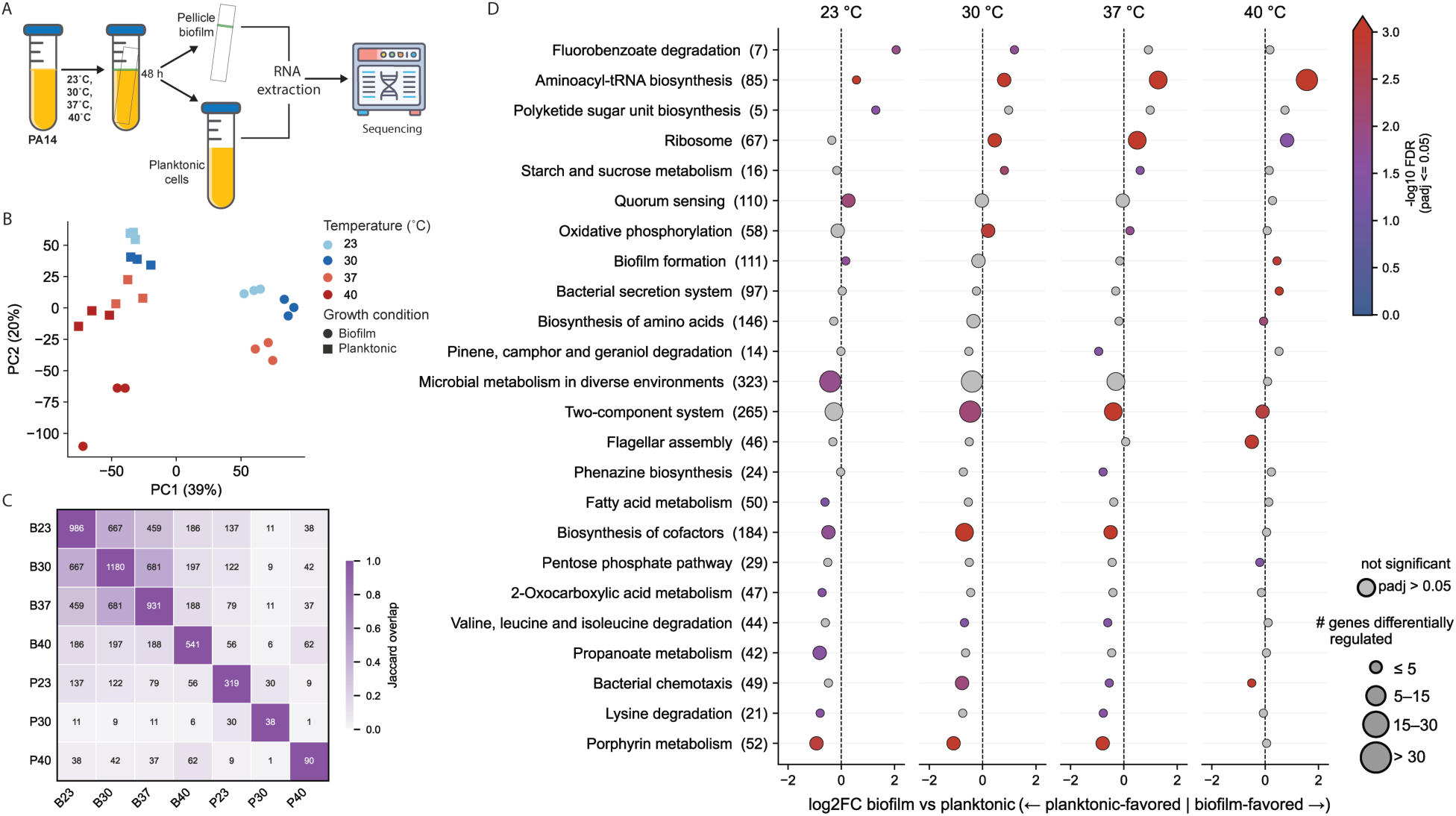
Temperature and growth state alter the transcriptomic landscape of *Pseudomonas aeruginosa* PA14. **(A)** Schematic of the RNA-seq experimental design. *P. aeruginosa* PA14 was grown overnight at 37°C and then sub-cultured in parallel at 23°C, 30°C, 37°C and 40°C. At 48 hours, RNA was extracted from biofilm and planktonic cells (n = 3 biological replicates per condition) and subjected to RNA sequencing. Differential gene expression analyses were performed using Rockhopper. **(B).** Principal component analysis (PCA) of transcriptomes from biofilm and planktonic *P. aeruginosa* PA14 populations grown at 23°C, 30°C, 37°C, and 40°C. Each point represents an independent biological replicate; colors indicate temperature and symbols denote growth state. **(C)** Pairwise overlap between the set of genes differentially expressed relative to planktonic growth associated cells at 37°C, for biofilm (B) and planktonic (P) cells at 23, 30, 37 and 40°C. Diagonal values give the number of genes in each set; off diagonal values give the number of genes shared between the two conditions. Color indicates the Jaccard overlap index (intersection/union), which is 1 on the diagonal. **(D)** KEGG pathway enrichment for biofilm versus planktonic comparison at 23, 30, 37 and 40°C. Panel represents the four temperatures. The x-axis is the mean log2 fold change (biofilm/planktonic), negative values indicate planktonic-favored expression, and positive values represent biofilm-favored expression. Dot size denotes the number of differentially regulated genes within the pathway and color denotes significance (-log10 FDR); grey dots are non-significant. Numbers in parentheses give the total number of genes annotated to each pathway.

To assess how growth state and temperature jointly shape the transcriptome, we compared each condition with 37°C planktonic growth associated cells, the optimal laboratory growth temperature for this bacterium, as the reference condition, and examined the pairwise overlap between the resulting gene sets **(Fig. 1C)**. Biofilm associated cells (B) showed a far broader response than planktonic cells (P), with 966, 1180, 931 and 541 differentially expressed genes at 23, 30, 37 and 40°C, respectively, compared with 319, 38 and 90 genes in planktonic associated cells at 23, 30 and 40°C. Within the biofilm, the three lower temperatures shared a substantial core (667 genes between B23 and B30, 681 between B30 and B37, and 459 between B23 and B37), whereas B40 overlapped to a lesser extent with the remaining biofilm conditions (187-197 genes), suggesting a distinct transcriptional program at this temperature. Overlap between biofilm and planktonic sets was consistently limited (<137 genes), and the small gene sets at P30 and P40 indicate that, in planktonic cells, shifting from 37°C to 30 or 40°C perturbs expression only marginally. Collectively, these results indicate that growth state is the principal determinant of the transcriptional response, with temperature acting as a strong secondary influence within the biofilm and a comparatively weak one in planktonic cells.

To gain further insight into temperature- and growth state-dependent responses, we performed KEGG pathway enrichment analysis on the genes differentially regulated between biofilm and planktonic cells at each temperature, which identified both conserved and temperature-specific responses **(Fig. 1D)**. Aminoacyl-tRNA biosynthesis was the only pathway significantly biofilm-favored at all the four temperatures, with effect size increasing with temperature. Regulatory and signaling pathways showed temperature-dependent responses: two-component systems were significantly planktonic-favored at 30, 37 and 40°C, whereas quorum sensing was planktonic-favored only at 23°C. At 40°C, flagellar assembly and bacterial chemotaxis were strongly planktonic-favored, while biofilm formation and bacterial secretion system were biofilm-favored. In contrast, porphyrin metabolism and biosynthesis of cofactors were consistently planktonic-favored at 23-37°C, but not at 40°C. Overall, differential enrichment was concentrated within specific regulatory, motility and biofilm associated pathways rather than broad metabolic functions. The temperature-dependent behavior of these pathways, several of which contribute to virulence, prompted us to examine virulence-associated genes directly.

### Virulence-associated secondary metabolite pathways exhibit temperature-dependent regulation

We next turned to virulence factors **(Table S1)** that were differentially regulated at environmental versus host temperatures in both biofilm and planktonic growth states. To this end, we calculated the log2 fold change in gene expression of virulence factor for biofilm and planktonic associated cells, comparing environmental with host temperature, and additionally compared biofilm versus planktonic cells at each of the four temperatures. Two signatures emerged consistently across every comparison and in both growth states: the T6SS associated genes *hcpA–D* were elevated at the lower temperature, whereas *exsA*, the master regulator of T3SS, was elevated at 37°C and 40°C **(Fig. 2A and B, Fig. S2A)**. This temperature-dependent induction of T3SS at host temperature is consistent with previous reports (18). T6SS mediates contact-dependent killing of competitors through a phage-like puncturing mechanism delivering effectors such as Hcp, shown to be thermoregulated in *P. aeruginosa* and *V. cholerae* (26) (27) (28, 29). T3SS on the other hand promotes host cell lysis and pathogenesis through effectors like Exo proteins and master regulator *exsA* (30, 31).

**Fig. 2.**
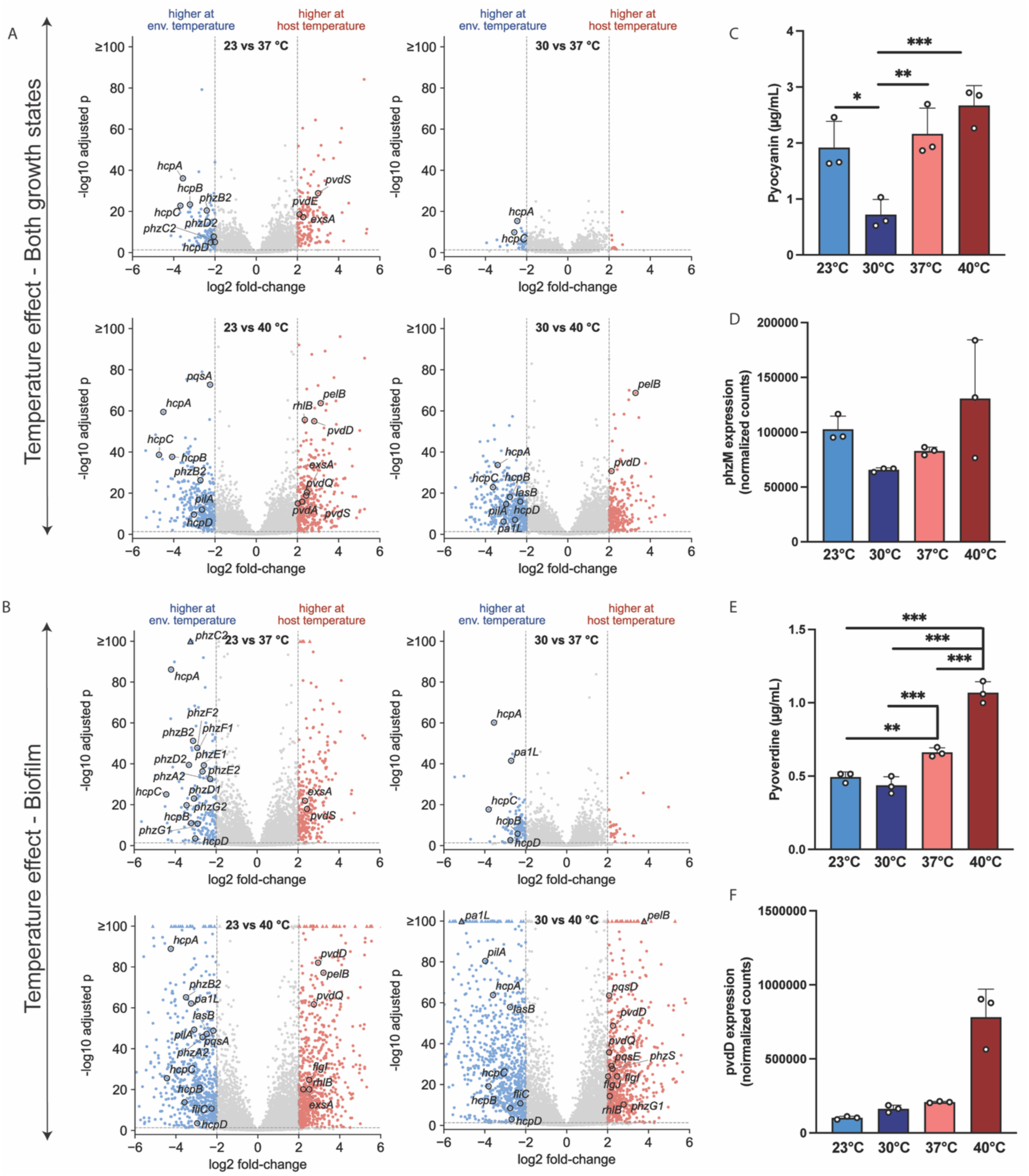
Effect of temperature and growth state on virulence factors. (A, **B)** Volcano plots of genes differentially expressed at host versus environmental temperature, shown for all cells irrespective of growth state **(A)** and for biofilm-associated cells **(B)**. Genes passing the significance threshold (q-value < 0.05, |log2 FC| ≥ 2) are plotted; blue and red dots denote significantly upregulated genes in environment versus host temperatures, respectively. Genes of interest are annotated. **(C)** Pyocyanin production by PA14 WT grown statically in tubes for 48 h at 23, 30, 37, or 40 °C, extracted from culture supernatants as described in Experimental Procedures. **(D)** Normalized expression of *phzM* in the biofilm growth state. **(E)** Pyoverdine production, measured spectrophotometrically (A405) and normalized to the OD600 of the corresponding culture. **(F)** Normalized expression of *pvdD* in the biofilm growth state. Values represent means ± SD of three independent experiments. Statistical significance was determined by two-tailed unpaired *t-*test: \**P* < 0.05, \*\**P* < 0.01, \*\*\**P* < 0.001.

Another striking temperature-dependent difference emerged in the expression of genes associated with pyocyanin and pyoverdine, two major virulence factors that enable *P. aeruginosa* to compete and survive under stressful conditions. Pyocyanin is an active phenazine and a well-established redox-active toxin. Once secreted, it interferes with a range of cellular functions by inducing oxidative stress and cell lysis, and it also contributes to biofilm formation by significantly increasing eDNA release (32, 33). In soil, pyocyanin further increases phosphorus bioavailability by liberating it from iron minerals through redox cycling (34, 35). Pyoverdine on the other hand is a siderophore that sequesters iron, making it accessible to this pathogen while preventing its usage by other competing microbes (36, 37). Our transcriptomic data revealed distinct regulatory patterns for the genes associated with these two virulence factors. For pyocyanin, taking *phzM* as the gene of interest, expression was highest at 40°C and lowest at 30°C, whereas for pyoverdine, taking *pvdD* as the gene of interest, expression increased steadily with temperature **(Fig. 2D and F)**. To determine whether these transcriptional trends were reflected at the phenotypic level, we performed biochemical analyses by extracting and quantifying both pigments at each temperature using established protocols (38, 39). Because these pigments cannot be extracted from the biofilm state, we used supernatants from static cultures grown for 48 h at each of the four temperatures. Pyocyanin production was lowest in cells grown at 30°C, comparable at 23°C and 37°C, and highest at 40°C **(Fig. 2C)**, a temperature-dependent pattern also reported previously (40). Pyoverdine production increased with temperature, although cultures grown at 23°C and 30°C accumulated nearly identical amounts **(Fig. 2E)**. Together, the RNA-seq and biochemical results, both highlight the role of temperature as a key regulator of these two important virulence factors.

We next examined the virulence-associated genes that were differentially expressed between biofilm and planktonic states, independent of temperature (**Fig. S2B**). Several genes linked to quorum sensing, biofilm development, and surface attachment were enriched in biofilms. The quorum-sensing regulator *rhlR* (41) showed higher expression in biofilms. The phenazine biosynthesis gene *phzA2* and *phzB*2 were consistently upregulated in biofilms across all four temperatures. Similarly, *pa1L* and *lasB*, encoding the lectin and elastase B, respectively, showed increased expression in the biofilm state at all temperatures. Both genes are quorum sensing–dependent and have previously been implicated in biofilm formation and maturation (42, 43). Finally, *pilA*, which encodes the major structural subunit of type IV pili (44), also showed increased expression in the biofilm state.

Given our previous observation that the PA14 wild-type strain forms its most robust biofilms at 23°C, we next investigated the contribution of candidate genes identified through our transcriptomic analysis to biofilm formation using corresponding mutants from the PA14 transposon mutant library (45). Δ*lasB* and Δ*pa1L* mutants resulted in significantly altered biofilm biomass at all four temperatures. In contrast, the Δ*exsA* mutant showed no significant difference in biofilm formation relative to the wild type. The Δ*hcpA* mutant exhibited a significant biofilm defect at 23°C and 30°C, whereas Δ*pvdD* displayed significant differences at all temperatures except 23°C. Similarly, Δ*phzM* as well as Δ*phzA2* mutant showed significantly altered biofilm biomass at 30°C and 37°C (**Fig. S2C**).

Overall, our transcriptomic analysis as well as the biochemical assays, indicates that *P. aeruginosa* virulence gene expression is shaped by two largely separable inputs: growth state, which governs a consistent set of quorum sensing–associated factors, and temperature, which restructures a distinct subset of the virulence transcriptome, most prominently at febrile temperature.

### EPS matrix-targeting enzymatic and chemical treatments differentially disrupt environment- and host temperature-associated biofilms

Our previous work demonstrated temperature-dependent differences in biofilm matrix composition and revealed global proteomic variation among biofilms formed at 23°C and 37°C (21, 46). Consistent with these observations, PA14 colonies grown for 48 h on Congo red/Coomassie blue agar displayed distinct morphologies across all four temperatures. This assay uses Congo red to detect matrix polysaccharides and Coomassie blue to detect matrix-associated proteins. Colonies grown at 23°C exhibited greater apparent Congo red staining and reduced Coomassie blue staining compared to colonies grown at the higher temperatures, suggesting temperature-dependent differences in matrix composition **(Fig. 3A)**.

**Fig 3.**
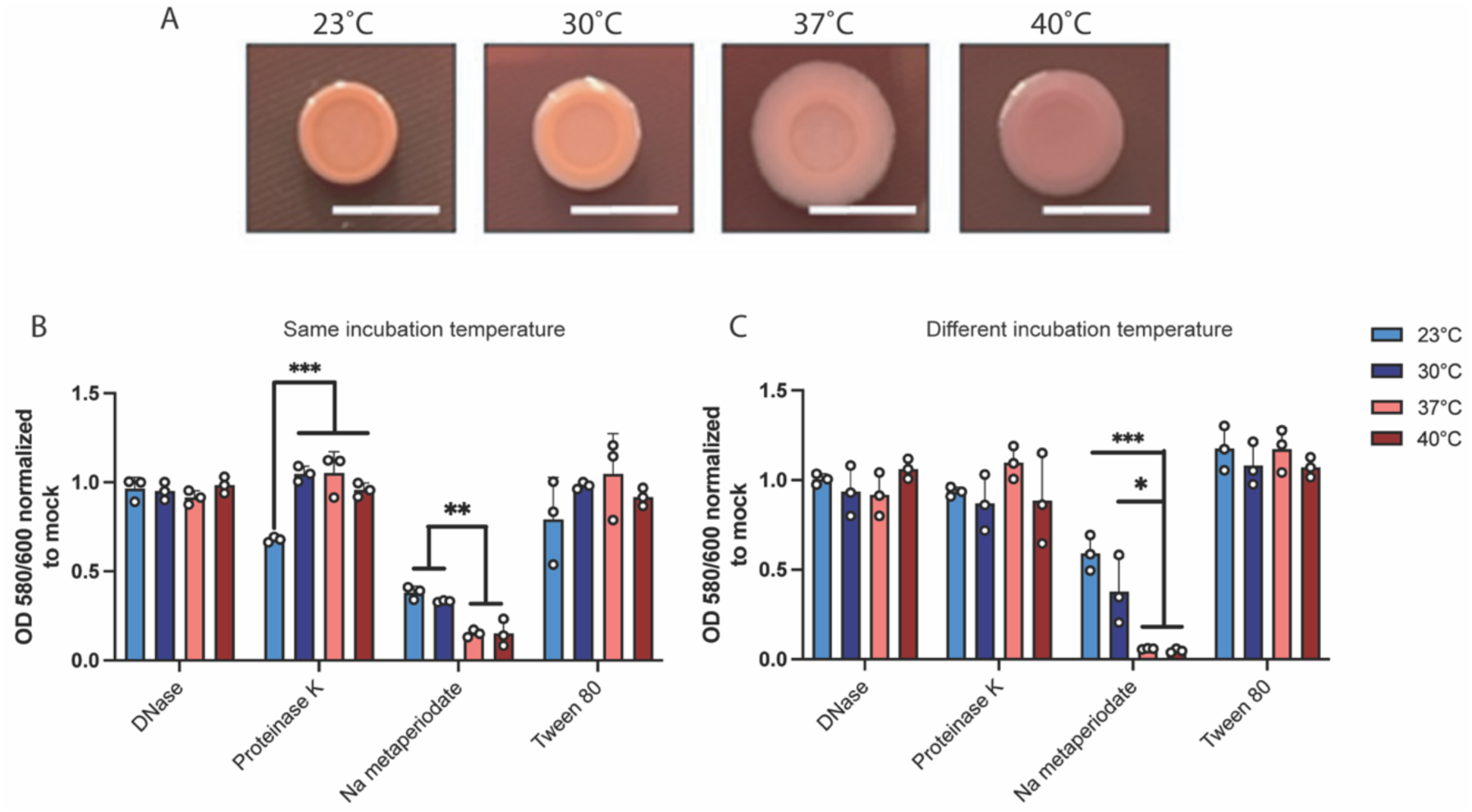
Effect of matrix degrading enzymes on *P. aeruginosa* biofilms. **(A)** Congo red binding assay reveals temperature specific difference in colony morphology. Extracellular matrix production by the wildtype was evaluated on tryptone agar plates containing Congo Red and Coomassie brilliant blue G after incubation at the four temperatures for 48 h. Representative images of the colony morphologies of PA14 are shown. Scale bar: 1 cm. Biofilms were grown for 48 hours at 23°C, 30°C, 37°C, and 40°C, after which they were subjected to different treatments. **(B)** Biofilm biomass of *P. aeruginosa* cells subjected to the same incubation temperature during treatment (32°C). **(C)**. Biofilm biomass of *P. aeruginosa* cells subjected to their respective growth temperature during treatment. Bars represent the mean of biological replicates performed on different days. The mean of each biological replicate was based on six technical replicates. Error bars represent the standard error of mean of the biological replicates. Unpaired t-test (two-tailed) was used to measure statistical significance. * *P*< 0.05, \*\**P* < 0.01, and \*\*\**P* < 0.001.

To exploit the observed differences in extracellular polymeric substance (EPS) components, we selectively targeted major matrix constituents, including proteins, carbohydrates, lipids, and extracellular DNA (eDNA), using established enzymatic and chemical treatments. DNase I and Proteinase K were used to degrade eDNA and protein components of the matrix, respectively (47, 48), while sodium metaperiodate was used to disrupt polysaccharide components (49). Tween 80, a nonionic surfactant with known antimicrobial and biofilm-disrupting properties, was used to target matrix-associated lipids and membrane interactions (50, 51). Biofilms were established at each of the four temperatures for 48 h and subsequently exposed to the respective treatments for 1 h. In one setup, all plates were incubated at a common temperature of 32°C during treatment **(Fig. 3B)**, whereas in the second setup, plates were maintained at their respective biofilm growth temperatures during treatment **(Fig. 3C)**.

Sodium metaperiodate treatment resulted in a significant reduction in biofilm biomass specifically in biofilms formed at host-associated temperatures. This reduction was observed under both treatment setups, indicating an increased dependence on polysaccharide matrix components at higher temperatures. In contrast, Proteinase K treatment significantly reduced biomass only in biofilms formed at 23°C when treated at 32°C, suggesting a greater contribution of proteinaceous matrix components under environmental growth conditions. DNase I and Tween 80 treatments produced comparatively limited effects on biofilm biomass across temperatures. Collectively, these findings support the presence of distinct matrix architectures in biofilms formed at environmental versus host-associated temperatures, with host-temperature biofilms exhibiting a greater reliance on polysaccharides and environmental temperature biofilms showing increased sensitivity to protein disruption.

### Environment- and host temperature-associated biofilms exhibit distinct stress tolerance profiles

We next examined whether biofilms formed at different temperatures differed in their susceptibility to widely used environmental and chemical stressors. Our previous study demonstrated temperature-dependent differences in biofilm architecture and physiology (46), leading us to hypothesize that biofilms formed at environmental (23°C and 30°C) and host-associated (37°C and 40°C) temperatures would exhibit distinct stress tolerance profiles. To test this hypothesis, 48 h biofilms were exposed to antibiotics, pH stress, osmotic stress, methanol, or hydrogen peroxide **(Fig. 4A)**. Following a 4 h treatment period, biofilms were sonicated, and biofilm-associated cells were serially diluted and plated to determine the number of viable cells remaining after exposure.

**Fig 4.**
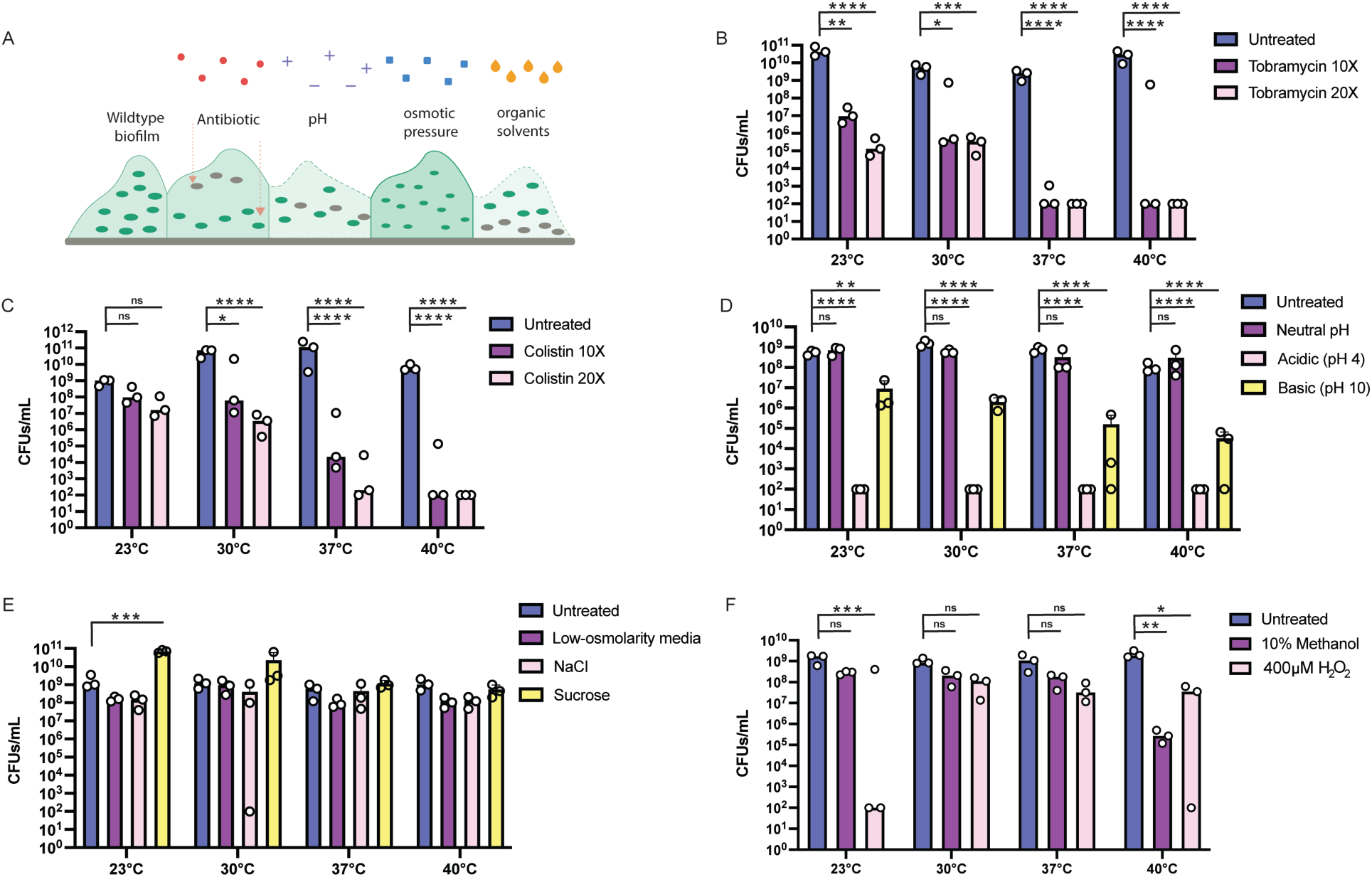
Stressor susceptibility profile of *P. aeruginosa* biofilms grown at four temperatures. **(A)** Representative schematic of the biofilm stress assay, each stress condition was assayed independently. In the schematic, green cells: viable cells, grey cells: stressed or lysed cells, shading and border thickness depict biofilm matrix density. Biofilms were grown for 48 h at each of the four temperatures. Following biofilm formation, planktonic cells were removed by aspiration, and biofilms were exposed independently to the indicated stressors for 4 h. Biofilm-associated cells were then recovered by sonication, serially diluted, and plated on Luria agar plates for overnight incubation at 37°C. Colony-forming units (CFU) were subsequently quantified. **(B and C)** Bar graph representing CFU/mL for *P. aeruginosa* biofilms following treatment with the antibiotics tobramycin and colistin. Tobramycin and colistin were applied at their respective minimum inhibitory concentrations (MICs) of 2 μg/mL and 1 μg/mL**. (D)** Bar graph representing CFU/mL for *P. aeruginosa* biofilms following exposure to pH stress conditions (pH 4, 7, and 10). **(E)** Bar graph representing CFU/mL for *P. aeruginosa* biofilms following exposure to osmotic stress conditions, including low-osmolarity medium, sodium chloride and sucrose. **(F)** Bar graph representing CFU/mL for *P. aeruginosa* biofilm following exposure to oxidative and solvent stress, including 400 µM H2O2 and 10% methanol, respectively. Error bars represent the mean of three independent biological replicates performed on different days. The mean of each biological replicate was based on three technical replicates. Statistical significance was determined using a two-way ANOVA on log10 transformed CFU counts with Dunnett’s multiple-comparisons correction. * *P*< 0.05, \*\**P* < 0.01, \*\*\**P* < 0.001, and \*\*\*\**P* < 0.0001.

We first assessed susceptibility to tobramycin and colistin, two antibiotics commonly used to treat *P. aeruginosa* biofilm-associated infections. Tobramycin primarily targets metabolically active cells, whereas colistin is effective against subpopulations that include biofilm-associated persister cells (52, 53). Both antibiotics are frequently used in combination for the treatment of *P. aeruginosa* infections (54, 55). For both antibiotics, susceptibility increased with the temperature at which the biofilm was formed **(Fig. 4B, C)**. Tobramycin significantly reduced viable counts at every temperature tested, but the magnitude of killing was temperature dependent. At 23°C and 30°C, 10X and 20X tobramycin reduced CFUs by approximately 3-5 logs relative to untreated controls, while at 37°C and 40°C, both concentrations reduced viable counts to the limit of detection **(Fig. 4B)**. Colistin showed the same trend, but with a sharper transition between environmental and host-associated temperatures. Biofilm formed at 23°C were not affected by either concentration of colistin, while biofilms formed at 30°C were reduced by roughly 3-4 logs. In contrast, biofilms formed at 37°C and 40°C were reduced to at or near the limit of detection **(Fig. 4C)**.

We next examined the effects of pH stress on biofilms formed at environmental and host-associated temperatures. Biofilms were exposed to pH 4, 7, and 10 solutions, and viable cell counts were determined following treatment. Exposure to pH 4 as well as pH 10 reduced biofilm-associated viability at all temperatures tested **(Fig. 4D)**. Our results are consistent with the findings of some other groups showing a decrease in biofilm formation at a lower pH (56). In contrast, exposure to pH 7 had no significant effect on viable cell counts at any temperature.

Since cells in the biofilm can be impacted by osmotic stress differently in natural versus human body environment (57), we next evaluated the impact of osmotic stress on biofilms formed at different temperatures. To mimic hypoosmotic and hyperosmotic conditions, biofilms were exposed to a low-osmolarity medium, NaCl, or sucrose. Treatment with both the low-osmolarity medium and NaCl resulted in greater reductions in viable biofilm-associated cells at 23°C and 40°C compared with other temperatures tested, however, these differences were not statistically significant. In contrast, sucrose had little effect on biofilm viability across all temperatures. Notably, sucrose treatment was associated with increased viable cell counts in biofilms formed at 23°C **(Fig. 4E)**.

Finally, we assessed the effects of methanol and hydrogen peroxide on biofilm viability. Biofilms were treated with 10% methanol or 400 μM H₂O₂, and viable cell counts were determined following exposure. Both treatments have been studied previously for their role as an antibiofilm agent in both environment and host associated pathogens (58–61). Methanol treatment had no significant effect on biofilms formed at 23°C, 30°C, or 37°C. Biofilms formed at 40°C were the exception, showing a reduction of roughly four logs compared to untreated control **(Fig. 4F)**. In contrast, H₂O₂ treatment produced the largest decrease in viable cell counts in biofilms formed at 23°C, with no significant effect at 30°C and 37°C **(Fig. 4F)**. At 40°C, H₂O₂ treatment caused a modest but significant reduction of approximately 1.5 logs. These results demonstrate that susceptibility to methanol and oxidative stress varies according to the temperature at which biofilms are formed. Overall, biofilms formed at environmental and host-associated temperatures displayed distinct susceptibility profiles across the tested stressor conditions, indicating that growth temperature is an important determinant of biofilm stress tolerance.

## Discussion

*Pseudomonas aeruginosa* occupies a remarkable diversity of environmental and host-associated niches and must continuously adapt to fluctuations in temperature, nutrient availability, and physicochemical stress. While temperature is well recognized as an important environmental signal in bacterial pathogens, the extent to which it coordinates biofilm physiology and virulence in *P. aeruginosa* has remained less understood (62, 63). In this study, we demonstrate that temperature strongly remodels the *P. aeruginosa* transcriptome and that these changes extend beyond gene expression to influence virulence factor production, biofilm matrix composition, and stress tolerance. Collectively, our findings highlight the role of temperature as a key environmental cue that helps coordinate the transition of *P. aeruginosa* from environmental reservoirs to mammalian hosts.

One of the most striking observations from our transcriptomic analysis was the large-scale restructuring of gene expression across both temperature and growth state. Consistent with previous studies demonstrating extensive transcriptional differences between planktonic and biofilm-associated cells, growth state represented a major source of variation in the dataset. However, temperature independently contributed to substantial transcriptome remodeling, particularly in biofilms. The greatest transcriptional divergence occurred at 40°C, where biofilm-associated cells displayed a distinct expression profile relative to all other conditions. This finding suggests that febrile temperatures represent more than a simple extension of host-associated growth conditions and may trigger specialized adaptive responses. Fever constitutes a common host defense mechanism, and the extensive transcriptomic changes observed at 40°C imply that *P. aeruginosa* actively senses and responds to this signal rather than merely tolerating thermal stress.

A major finding of this temperature-dependent transcriptional remodeling was the differential regulation of virulence-associated pathways. Among the most consistent signatures identified were the reciprocal expression patterns of T6SS- and T3SS-associated genes. The elevated expression of *hcp* genes at environmental temperatures and increased expression of the T3SS regulator *exsA* at host temperatures suggest a temperature-mediated shift in ecological strategy. The T6SS is widely associated with interbacterial competition and environmental fitness (64), whereas the T3SS primarily facilitates interactions with eukaryotic hosts (65). Enhanced expression of T6SS-associated components at lower temperatures may therefore provide a competitive advantage in polymicrobial environmental communities, while induction of T3SS-associated genes at 37°C and 40°C would favor host colonization and pathogenesis. These observations support the notion that temperature serves as a predictive environmental signal that allows *P. aeruginosa* to anticipate the selective pressures associated with environmental versus host-associated habitats.

Temperature-dependent regulation was also evident in the production of the secondary metabolites pyocyanin and pyoverdine. Transcriptomic analyses and biochemical measurements showed a strong induction of pyocyanin synthesis at 40°C and a progressive increase in pyoverdine production with increasing temperature. Pyocyanin contributes to oxidative stress, host tissue damage, and biofilm development through eDNA release (33, 66), whereas pyoverdine functions both as an iron-scavenging molecule and as a regulator of additional virulence traits (36). Enhanced production of these factors at host-associated temperatures is consistent with the increased importance of iron acquisition and host manipulation during infection. Interestingly, pyocyanin displayed maximal production specifically at febrile temperature rather than at 37°C, further suggesting that 40°C induces a unique physiological state. Since fever is associated with immune activation and altered nutrient availability within the host, increased production of these secondary metabolites may provide a mechanism for maintaining fitness during infection under hostile host conditions.

In contrast to temperature-regulated virulence factors, a distinct set of genes responded to growth state rather than temperature. This suggests that *P. aeruginosa* does not regulate its virulence genes on and off as one unit, temperature controls some, while the shift from free-floating planktonic to surface-attached biofilm growth state controls others. Several quorum sensing-dependent genes, including *rhlR*, *lasB* and *pa1L*, displayed consistent enrichment in biofilms regardless of temperature. These observations reinforce the importance of growth state as a distinct regulatory input governing virulence gene expression. Biofilms are characterized by dense cellular communities in which quorum sensing signaling is highly active, and the temperature-independent expression of these genes likely reflects their fundamental roles in biofilm establishment and maintenance (67). The biofilm phenotypes observed in the corresponding mutants further support this interpretation, as disruption of *lasB* and *pa1L* altered biofilm development across all temperatures tested. *exsA* known to be highly expressed in host associated temperature, was not required for biofilm formation at any temperature, indicating that type III secretion is largely uncoupled form the surface-attached state under our conditions. Other genes showed temperature-restricted phenotypes. Loss of *hcpA* produced a biofilm defect at low temperatures, whereas *pvdD* was dispensable at 23°C but required at all other temperatures. T6SS and pyoverdine-mediated iron acquisition thus contribute under different physiological regimes, and the relative importance of individual determinants shifts with the thermal environment even when their transcription does not.

We also identified growth state-dependent regulation within the phenazine biosynthetic pathway. *phzA2* and *phzB2* were enriched in biofilms, consistent with the established role of phenazines as extracellular electron shuttles that support metabolism in oxygen-limited biofilm interiors and influence biofilm structure. Yet mutant phenotypes were restricted to a subset of conditions: loss of *phzA2* altered biofilm biomass at 30°C and 37°C but not at 23°C or 40°C, and disruption of *phzM* showed a similar pattern. Thus, biofilm-associated expression did not translate into equal requirement across all temperatures, suggesting that the contribution of phenazines depends on the physiological context in which they are produced. One explanation is functional overlap between the two phenazine operons. Previous study has shown that phz2 is preferentially expressed in biofilms and that the two operons are differentially regulated across growth states (68), raising the possibility that temperature-dependent shifts in operon usage could mask the effects of phzA2 under some conditions. Together, these findings suggest that growth state and temperature regulate largely separable components of the virulence network. Whereas biofilm formation appears to rely on a conserved quorum sensing and phenazine-associated regulatory program, temperature modifies a distinct subset of virulence traits associated with environmental survival and host adaptation.

Our findings further indicate that temperature-dependent transcriptional responses are associated with changes in biofilm matrix organization. Previous work from our lab demonstrated that biofilm architecture and proteome composition vary substantially with temperature, and the matrix disruption experiments performed here provide functional evidence that these differences translate into altered matrix dependencies (21, 46). Surprisingly, sodium metaperiodate treatment caused a greater reduction in biofilm biomass at host-associated temperatures despite the higher total polysaccharide content previously detected in 23°C biofilms (46). This suggests that biofilm susceptibility to periodate is determined not simply by polysaccharide abundance, but by the contribution of specific polysaccharide components to matrix integrity. At higher temperatures, polysaccharides may function as critical structural elements required for biofilm cohesion, whereas the more abundant polysaccharides present at lower temperatures may play a less central role or be compensated by other matrix components, such as proteins and extracellular DNA. In contrast, biofilms formed at 23°C was more sensitive to protein degradation by Proteinase K, indicating a greater dependence on proteinaceous matrix components. Together, these observations are consistent with the altered Congo red and Coomassie blue staining patterns observed across temperatures and support the existence of distinct matrix architectures optimized for different environmental conditions. Such temperature-dependent matrix remodeling may allow *P. aeruginosa* to maximize biofilm stability and protection against the specific stresses encountered in environmental reservoirs versus host-associated niches.

Importantly, these matrix differences were accompanied by marked differences in stress tolerance. Our stress susceptibility assays revealed that biofilms formed at environmental and host-associated temperatures are physiologically distinct. Biofilms established at 23°C and 30°C exhibited greater tolerance to clinically relevant antibiotics than those formed at 37°C and 40°C. This observation is consistent with current models of biofilm-associated antimicrobial tolerance, which is largely physiological rather than genetically encoded. For aminoglycosides such as tobramycin, reduced efficacy can arise from interactions with negatively charged matrix components including extracellular DNA, Pel, and alginate, which impede antibiotic penetration into the biofilm interior. Colistin, which acts at the outer membrane, is similarly affected by electrostatic sequestration within the matrix. In addition, nutrient and oxygen gradients within mature biofilms promote the formation of slow-growing persister cell populations that are inherently less susceptible to antibiotic killing (69–71). The greater tolerance observed in environmental-temperature biofilms to both agents may therefore reflect the distinct matrix architecture identified in our EPS disruption experiments, although we did not directly measure antibiotic penetration and cannot exclude contributions from temperature-dependent differences in metabolic activity. Colistin was ineffective against biofilms formed at 23°C at both tested concentrations, suggesting limited activity against environmentally conditioned biofilms. Given that sublethal antimicrobial exposure promotes resistance evolution (72), biofilm growth history should therefore be considered in treatment selection.

The differential response to pH stress further supports the idea that biofilm physiology varies substantially with growth temperature. Acidic and alkaline conditions are known to affect biofilms through distinct mechanisms (56, 73). Low pH can alter cell surface charge and protonate extracellular matrix polymers, in some cases initially enhancing adhesion and matrix cohesion, whereas alkaline conditions frequently promote matrix destabilization and biofilm dispersal (74, 75). Neutral pH had no effect on viability at any temperature. In contrast exposure to pH 4 reduced viable cell count to the limit of detection at all four temperatures. Alkaline stress (pH 10) also significantly reduced viability, though less severely than pH 4. These findings are consistent with prior work on pH-dependent biofilm regulation in diverse bacterial species (56). One possible explanation for this temperature-dependent response is that pH modulates the activity, stability, and structural integrity of matrix-associated proteins. Because our previous work demonstrated substantial temperature-dependent differences in the biofilm proteome and matrix composition (21), distinct protein complements present at different temperatures could confer variable sensitivity to acidic and alkaline conditions. These observations emphasize that pH-based interventions may not exert uniform effects across all biofilm states and instead depend on the physiological conditions under which the biofilm was established.

Osmotic stress had comparatively little effect on biofilm viability. Neither low-osmolarity medium nor NaCl significantly reduced viable cell counts at any temperature tested. Sucrose likewise did not reduce viability, and at 23°C significantly increased viable cells compared to untreated biofilms. A similar trend was observed at 30°C, however, it was not significant. This resistance is consistent with the well-described capacity of *P. aeruginosa* to buffer changes in water activity through accumulation of compatible solutes (57, 76, 77). Exposure to elevated osmolarity can stimulate production of matrix polysaccharides while promoting accumulation of osmo-protectants such as glycine betaine, trehalose, and N-acetylglutaminylglutamine amide (78, 79). The divergent response to sucrose versus NaCl further suggests that the identity of the osmolyte, not osmolarity alone, shapes the outcome. The increase viable cell count with sucrose could be a result of either enhanced growth or improved disaggregation of the biofilm.

Responses to methanol and oxidative stress were strongly temperature-dependent, and interestingly, neither treatment significantly reduced viability in biofilms formed at 37°C. Methanol produced no significant reduction at 23°C and 30°C, and its largest effect at 40°C. H₂O₂ was most effective against biofilms formed at 23°C, reducing the viable cell counts by roughly seven logs, with smaller but significant reductions at 40°C. Unlike antibiotics, whose effectiveness is often influenced by diffusion barriers and metabolic heterogeneity, tolerance to organic solvents is more strongly associated with active cellular defense mechanisms, including resistance-nodulation-cell division (RND) efflux systems and membrane remodeling processes (80). In *P. aeruginosa*, MexAB-OprM and related efflux pumps play central roles in solvent resistance, while cis-to-trans isomerization of membrane fatty acids helps maintain membrane integrity in the presence of membrane-disrupting compounds (81). Enhance methanol susceptibility at 40°C may therefore reflect increased membrane fluidity at elevated temperature, which in combination with solvent stress, may further compromise membrane integrity. Oxidative stress tolerance similarly depends on inducible defenses, and the pronounced susceptibility of 23°C biofilms suggest these may be less fully engaged at environmental temperatures.

Taken together, our findings demonstrate that biofilms formed at environmental and host-associated temperatures possess distinct physiological, structural, and transcriptional states. Temperature-dependent differences in virulence factor expression, matrix composition, antimicrobial susceptibility, and stress tolerance indicate that biofilms formed under these conditions are not simply thermal variants of the same community. Rather, they represent alternative adaptive states optimized for survival in different ecological niches. Environmental-temperature biofilms appear to favor traits associated with persistence, interbacterial competition, whereas host-range temperatures promote the expression of factors that enhance host interaction and pathogenic potential.

These observations have important implications for both environmental control and clinical management of *P. aeruginosa*. Treatments that were effective against biofilms formed at one temperature often displayed substantially reduced efficacy against biofilms formed at another, despite involving the same bacterial strain. This finding highlights the importance of considering not only the genetic identity of a pathogen but also its environmental history when designing eradication strategies. The widespread use of standardized laboratory growth conditions (37°C) may therefore overlook physiologically relevant states that influence biofilm resilience and treatment outcomes in natural, industrial, and host-associated settings. More broadly, our findings support a model in which temperature functions not merely as a physical parameter or stressor, but as a predictive environmental cue that enables *P. aeruginosa* to anticipate and prepare for changes associated with distinct ecological niches. Rather than mounting a transient response to thermal fluctuations*, P. aeruginosa* appears to use temperature information to coordinate extensive reprogramming of its physiology, including virulence factor production, biofilm matrix architecture, and stress adaptation. This thermal responsiveness likely facilitates successful transitions between environmental reservoirs and mammalian hosts, allowing it to optimize traits required for persistence, competition, and infection before encountering the selective pressures of a new habitat. Our findings therefore position temperature as a central signal integrating environmental sensing with lifestyle adaptation in this versatile opportunistic pathogen. Future studies aimed at identifying the thermosensory mechanisms and regulatory networks underlying these responses will be critical for understanding how temperature intersects with established virulence, quorum-sensing, and biofilm regulatory pathways. Such insights may reveal novel strategies to manipulate temperature-dependent physiological states and improve the control of *P. aeruginosa* biofilms in both environmental and clinical settings.

## Materials and Methods

### Bacterial strains, media, and growth conditions

*P. aeruginosa* strain UCBPP-PA14, a highly virulent strain of *P. aeruginosa* originally isolated from a wound infection, was used in all experiments unless otherwise stated (82). Isogenic mutants of PA14, namely, PA14/MrT7::*PA14_33650* (pvdD), PA14/MrT7::*PA14_09490 (phzM),* PA14/MrT7::*PA14_39970* (*phzA2*), PA14/MrT7::*PA14_48890* (*hcpA*), PA14/MrT7:: *PA14_42390* (*exsA*), PA14/MrT7:: *PA14_31290* (*pa1L*), and PA14/MrT7:: *PA14_16250* (*lasB*), carrying the Mariner transposon *MAR2Tx7* (45), were used to examine biofilm formation at the four temperatures. Strains were routinely grown overnight and maintained at 37°C in Luria-Bertani (LB) broth. Gentamicin was added at 15 ug/ml to maintain the transposon in the mutants.

### Congo red binding assay

To phenotypically assess the different components of the EPS matrix, a colony morphology assay was performed as described previously (83). Extracellular matrix production by *P. aeruginosa* PA14 was evaluated on tryptone agar plates containing Congo Red and Coomassie brilliant blue G after incubation at 23°C, 30°C, 37°C and 40°C for 24, 48 and 72 hours. Five microliters of overnight precultures were spotted on Petri plates containing 20 ml of the assay medium (1% tryptone, Congo red dye (40 µg/ml), Coomassie brilliant blue G dye (20 µg/ml) and 1% agar). Colonies were grown at 23°C, 30°C, 37°C and 40°C for 48 hours. Images of the colonies were taken daily using the Nikon camera.

### Bacterial RNA extraction

A 20ul of the overnight grown culture of PA14 was added into a fresh 20ml LB broth, in a 50 ml falcon along with a microscope glass slide. This was left at static conditions at 23°C, 30°C, 37°C and 40°C for 48 hours. Three biological replicates were processed for the WT at each temperature. Next, the biofilm from the glass slides was scraped off using a tip and 700 ul of Qiazol lysis reagent. For disrupting or lysing bacterial cells, the mixture was added onto tubes with beads (Tough Microorganism Lysing Mix” (RNase+DNase free, size-2mL X 0.5mm Verre)) and lysed in a VWR Bead Mill Homogenizer at 6500 rpm for 1 min (two rounds). For planktonic cells, the 48 hours grown culture was centrifuged at 5000 rpm for 10 minutes, followed by discarding the supernatant and resuspending the pellet in 700 ul of Qiazol lysis reagent. RNA was then extracted using the RNeasy minikit (Qiagen) according to the manufacturer’s recommendations, and the RNA solution was digested with the RNase-free DNase set (Qiagen), followed by on-column DNase digestion to eliminate any remaining traces of genomic DNA. The purified RNA was quantified using a Take3 plate reader (Synergy H1 microplate reader, Biotek). RNA samples with 1.8 to 2.2 ratio of absorbance at 260/280 nm were kept for further analysis. The samples were then sent to Genewiz for library prep and Illumina HiSeq. Only samples with an RNA integration number (RIN) greater than 8.0 were used for cDNA library preparation.

### Analysis of the RNA-seq data

RNA-seq data was analyzed using Rockhopper software implementing reference-based transcript assembly with UCBPP-PA14 as a reference genome followed by calculating the fold change for the transcripts at each temperature (84). The full accession number of the UCBPP-PA14 annotation that was used is NC_008463. Each data set was normalized by upper quartile normalization, and then transcript abundance was quantified using reads assigned per the kilobase of target per million mapped reads (RPKM) normalization method.

Differential expression was determined using DESeq2, either by pairwise comparison between temperatures within each growth condition or using additive effects of temperature and growth condition as indicated. The selection criteria for differential expression required genes to have a fold change of ≥2 and a *q* value of ≤0.05 to be considered significant. The *q* value was obtained by adjusting the *P-*value using the Benjamini-Hochberg procedure (85). KEGG pathway enrichment was tested per temperature with a two-sided Mann-Whitney U test comparing each pathway’s fold-change values to those of all other tested genes, and per cluster with a hypergeometric over-representation test against the annotated gene background. *P* values were Benjamini-Hochberg-adjusted within each temperature or cluster (85).

### Microtiter biofilm formation assay

To quantify and study biofilm formation at each temperature we used the previously established microtiter biofilm formation assay(86). Wild-type *Pseudomonas aeruginosa* PA14 and the mutant strain were grown overnight in a 96 well round bottom plate in 150 microliters LB broth with shaking at 220 rpm and 37°C overnight. Next day, five microliters of the overnight grown culture were transferred to a fresh 96 well plate with 145 μL of LB media. This was done for 3 replicates at 3 different days. The plates were incubated for 48 hours at 23°C, 30°C, 37°C and 40°C. An absorbance at 600 nm wavelength was taken after 48 hours using Synergy Hi5 Microplate Reader, Biotek. Planktonic cells were then aspirated out, and the remaining biofilm was washed 3 times with 300 microliters of PBS. This step helps remove unattached cells and media components that can be stained in the next step and significantly lowers background staining. Next, 200 microliters of 100% ethanol was added to the wells and incubated for 15 minutes. The ethanol was then aspirated out completely and the plates are flipped upside down and left for drying. After the ethanol dried, 200 ul of a 1% solution of crystal violet (CV) was added to each well of the microtiter plate. The microtiter plate was then incubated for 15 minutes at room temperature followed by rinsing it 3-4 times with water by submerging the plate in a tub of water, and blot vigorously on a stack of paper towels to get rid of all the excess water. The microtiter plate was then left to dry for 1-2 hours. Finally, 150 microliters of 30% acetic acid solution was added to each well of the microtiter plate to solubilize the crystal violet. After an hour incubation at room temperature, absorbance was taken at 580 nm. Using the biomass baseline, this reading was quantified and analyzed to produce readable data.

### Pyocyanin quantification

Five-ml samples of the supernatant fractions of cultures of PA14 grown in LB for 48 hrs at the four temperatures were isolated, mixed with chloroform (1:1), and the lower layer was separated and mixed with two ml of 0.2 N HCl. The pyocyanin-rich organic upper layer (pink in color) was then extracted, and the absorbance at 520 nm was determined. Values were normalized by dividing by the OD_600_ of each culture prior application of the following formula. The amount of pyocyanin, in μg/ml, was calculated using the following formula: OD_520_ × 17.072 = μg of pyocyanin/ml(38).

### Pyoverdine assay

Pyoverdine levels within the supernatant of PA14 cultures grown in LB at the four temperatures for 48 hrs. were determined by measuring the *A*_405_ values of each sample(39). Values were adjusted by dividing *A*_405_ readings by the corresponding OD_600_ values of the culture.

### Matrix perturbation Assay

*Pseudomonas aeruginosa* was streaked out and grown overnight; the next day individual colonies were transferred to a new culture and grown in a shaking incubator overnight at 37°C. The overnight grown culture was next transferred to 96-wellplate and allowed to form a pellicle biofilm for 48 hours at 23°C, 30°C, 37°C, and 40°C in a static incubator. After this growth period, OD 600 was taken for all the plates using the Synergy H5 plate reader. The planktonic cells were then aspirated out and the biofilms remaining on the walls of the wells were treated with various enzymes and reagents for one hour at their respective temperatures and another set of plates were incubated at a constant temperature of 32°C. DNase, proteinase K, Na metaperiodate and Tween 80 was used for treating the biofilms. CV assay was then performed (as mentioned above). Absorbance was taken at 580 nm. Biomass was normalized to the buffer which were used as a control for each individual treatment. Using the biomass baseline, this reading was quantified and analyzed to produce readable data.

### Stressor Assay

*Pseudomonas aeruginosa* was streaked out and grown overnight; the next day individual colonies were transferred to a new culture and grown in a shaking incubator overnight at 37°C. The overnight grown culture was next transferred to tubes and allowed to form a pellicle biofilm for 48 hours at 23°C, 30°C, 37°C, and 40°C in a static incubator. After this growth period, the liquid culture, which mostly comprised of planktonic cells, was asphyxiated out and the biofilms remaining on the sides of the tubes were treated with various stressors for four hours at their respective temperatures. After 4 hours of incubation, the biofilm was sonicated, at 20% amplitude for 30 seconds, and the biofilm associated cells were serially diluted (10-folds) with the resulting cultures to determine the number of viable cells remaining after stressor treatment. The plates were then incubated overnight at 37°C incubator. Next day, colonies are counted from each plate to determine the colony forming units (CFU/mL) for each stressor treatment.

Antibiotic concentration used in this study were, for Tobramycin: MIC-2 μg/mL (Tobramycin T4014-500MG Sigma-Aldrich) and for Colistin: MIC-1 μg/mL (Colistin Sulfate C4461-1G Sigma-Aldrich)(87). For the osmolarity stressor assay a low osmolarity medium (40 mOsm/L) was used: K2HPO4 (1 mM), (NH4)2SO4 (1.5 mM), MgCl2 (0.08 mM), casein hydrolysate (4 g/L), trace metals (1 ml/L), and glucose (0.3 mM)(88). Osmolarity stress was induced by treating a 300mM sucrose (300 mOsm/L) (Millipore Sigma, S0389) or 1 M NaCl (2 Osm/L) (Fisher chemical, S271 –500) solution.

## Statistical analyses

Statistical analyses were performed using GraphPad Prism 9.0 (GraphPad Software, Inc., San Diego, CA). Unpaired t-test (two tailed) and two-way ANOVA was used to calculate the statistical significance.

## Data Availability

The raw RNA-seq data sets generated during this study are available through NCBI’s BioProject database under accession number PRJNA664520 and PRJNA1525626. The authors declare that all other relevant data supporting the claims of the paper are available either in the main text or Supplementary files.

## Acknowledgement

This work was supported by NIH/NIGMS grant R15GM128072 and NIH/NIAID grant R01AI173686 to CAW, and KB was supported by the Doctoral Dissertation Completion Fellowships granted from Texas Tech University Graduate School, Lubbock, TX

## References

1. Tolker-Nielsen T. 2014. Pseudomonas aeruginosa biofilm infections: From molecular biofilm biology to new treatment possibilities. APMIS 122:1–51.

2. Crone S, Vives-Flórez M, Kvich L, Saunders AM, Malone M, Nicolaisen MH, Martínez-García E, Rojas-Acosta C, Catalina Gomez-Puerto M, Calum H, Whiteley M, Kolter R, Bjarnsholt T. 2020. The environmental occurrence of Pseudomonas aeruginosa. APMIS 128:220–231.

3. Bowie KR, Luhung I, Burke TR, Roberts SC, Martinello RA, Gerstein M, Peccia J, Healy HG. 2026. Disinfection of hospital sink drains enriches pseudomonadota and efflux pump-mediated antibiotic resistance in reestablished biofilms. Nat Commun 17.

4. Veesenmeyer JL, Hauser AR, Lisboa T, Rello J. 2009. Pseudomonas aeruginosa virulence and therapy: evolving translational strategies. Critical care medicine 37:1777–1786.

5. Faure E, Kwong K, Nguyen D. 2018. Pseudomonas aeruginosa in Chronic Lung Infections: How to Adapt Within the Host? Frontiers in Immunology 9.

6. van ‘t Wout EFA, van Schadewijk A, van Boxtel R, Dalton LE, Clarke HJ, Tommassen J, Marciniak SJ, Hiemstra PS. 2015. Virulence Factors of Pseudomonas aeruginosa Induce Both the Unfolded Protein and Integrated Stress Responses in Airway Epithelial Cells. PLOS Pathogens 11:e1004946.

7. Newman JW, Floyd RV, Fothergill JL. 2017. The contribution of Pseudomonas aeruginosa virulence factors and host factors in the establishment of urinary tract infections. FEMS Microbiology Letters 364.

8. Høiby N, Bjarnsholt T, Givskov M, Molin S, Ciofu O. 2010. Antibiotic resistance of bacterial biofilms. International Journal of Antimicrobial Agents 35:322–332.

9. Chiang W-C, Nilsson M, Jensen PØ, Høiby N, Nielsen TE, Givskov M, Tolker-Nielsen T. 2013. Extracellular DNA shields against aminoglycosides in Pseudomonas aeruginosa biofilms. Antimicrobial agents and chemotherapy 57:2352–2361.

10. Costerton JW, Stewart PS, Greenberg EP. 1999. Bacterial biofilms: a common cause of persistent infections. Science 284:1318–22.

11. Kim M, Christley S, Khodarev NN, Fleming I, Huang Y, Chang E, Zaborina O, Alverdy JC. 2015. Pseudomonas aeruginosa wound infection involves activation of its iron acquisition system in response to fascial contact. The journal of trauma and acute care surgery 78:823–829.

12. Shapiro RS, Cowen LE. 2012. Thermal control of microbial development and virulence: molecular mechanisms of microbial temperature sensing. mBio 3.

13. Grosso-Becera MV, Servín-González L, Soberón-Chávez G. 2015. RNA structures are involved in the thermoregulation of bacterial virulence-associated traits. Trends Microbiol 23:509–18.

14. Grosso-Becerra MV, Croda-García G, Merino E, Servín-González L, Mojica-Espinosa R, Soberón-Chávez G. 2014. Regulation of Pseudomonas aeruginosa virulence factors by two novel RNA thermometers. Proc Natl Acad Sci U S A 111:15562–7.

15. Owings JP, Kuiper EG, Prezioso SM, Meisner J, Varga JJ, Zelinskaya N, Dammer EB, Duong DM, Seyfried NT, Albertí S, Conn GL, Goldberg JB. 2016. Pseudomonas aeruginosa EftM Is a Thermoregulated Methyltransferase. J Biol Chem 291:3280–90.

16. Almblad H, Randall TE, Liu F, Leblanc K, Groves RA, Kittichotirat W, Winsor GL, Fournier N, Au E, Groizeleau J, Rich JD, Lou Y, Granton E, Jennings LK, Singletary LA, Winstone TML, Good NM, Bumgarner RE, Hynes MF,…,Harrison JJ. 2021. Bacterial cyclic diguanylate signaling networks sense temperature. Nature Communications 12:1986.

17. Barbier M, Damron FH, Bielecki P, Suarez-Diez M, Puchalka J, Alberti S, Dos Santos VM, Goldberg JB. 2014. From the environment to the host: re-wiring of the transcriptome of Pseudomonas aeruginosa from 22 degrees C to 37 degrees C. PLoS One 9:e89941.

18. Wurtzel O, Yoder-Himes DR, Han K, Dandekar AA, Edelheit S, Greenberg EP, Sorek R, Lory S. 2012. The Single-Nucleotide Resolution Transcriptome of Pseudomonas aeruginosa Grown in Body Temperature. PLOS Pathogens 8:e1002945.

19. Robinson RE, Gebhardt MJ, Goldberg JB. 2026. The interplay between temperature and growth phase shapes the transcriptional landscape of *Pseudomonas aeruginosa*. J Bacteriol 208:e0038525.

20. Robinson RE, Robertson JK, Prezioso SM, Goldberg JB. 2025. Temperature controls LasR regulation of *piv* expression in *Pseudomonas aeruginosa*. mBio 16:e0054125.

21. Bisht K, Moore JL, Caprioli RM, Skaar EP, Wakeman CA. 2021. Impact of temperature-dependent phage expression on Pseudomonas aeruginosa biofilm formation. npj Biofilms and Microbiomes 7:22.

22. Kordes A, Preusse M, Willger SD, Braubach P, Jonigk D, Haverich A, Warnecke G, Häussler S. 2019. Genetically diverse Pseudomonas aeruginosa populations display similar transcriptomic profiles in a cystic fibrosis explanted lung. Nature communications 10:3397–3397.

23. Cornforth DM, Dees JL, Ibberson CB, Huse HK, Mathiesen IH, Kirketerp-Møller K, Wolcott RD, Rumbaugh KP, Bjarnsholt T, Whiteley M. 2018. Pseudomonas aeruginosa transcriptome during human infection. Proceedings of the National Academy of Sciences 115:E5125.

24. Gonzalez MR, Ducret V, Leoni S, Fleuchot B, Jafari P, Raffoul W, Applegate LA, Que Y-A, Perron K. 2018. Transcriptome Analysis of Pseudomonas aeruginosa Cultured in Human Burn Wound Exudates. Frontiers in cellular and infection microbiology 8:39–39.

25. Bielecki P, Komor U, Bielecka A, Müsken M, Puchałka J, Pletz MW, Ballmann M, Martins dos Santos VAP, Weiss S, Häussler S. 2013. Ex vivo transcriptional profiling reveals a common set of genes important for the adaptation of Pseudomonas aeruginosa to chronically infected host sites. Environmental Microbiology 15:570–587.

26. Yu K-W, Xue P, Fu Y, Yang L. 2021. T6SS Mediated Stress Responses for Bacterial Environmental Survival and Host Adaptation. International Journal of Molecular Sciences 22.

27. Chen L, Zou Y, She P, Wu Y. 2015. Composition, function, and regulation of T6SS in Pseudomonas aeruginosa. Microbiological Research 172:19–25.

28. Townsley L, Sison Mangus MP, Mehic S, Yildiz FH. 2016. Response of Vibrio cholerae to Low-Temperature Shifts: CspV Regulation of Type VI Secretion, Biofilm Formation, and Association with Zooplankton. Applied and Environmental Microbiology 82:4441.

29. Allsopp LP, Wood TE, Howard SA, Maggiorelli F, Nolan LM, Wettstadt S, Filloux A. 2017. RsmA and AmrZ orchestrate the assembly of all three type VI secretion systems in Pseudomonas aeruginosa. Proceedings of the National Academy of Sciences 114:7707.

30. Hauser AR. 2009. The type III secretion system of Pseudomonas aeruginosa: infection by injection. Nature reviews Microbiology 7:654–665.

31. Galle M, Carpentier I, Beyaert R. 2012. Structure and function of the Type III secretion system of Pseudomonas aeruginosa. Current protein & peptide science 13:831–842.

32. Hall S, McDermott C, Anoopkumar-Dukie S, McFarland AJ, Forbes A, Perkins AV, Davey AK, Chess-Williams R, Kiefel MJ, Arora D, Grant GD. 2016. Cellular effects of pyocyanin, a secreted virulence factor of Pseudomonas aeruginosa. Toxins 8:1–14.

33. Das T, Manefield M. 2012. Pyocyanin Promotes Extracellular DNA Release in Pseudomonas aeruginosa. PLoS ONE 7.

34. Hernandez ME, Kappler A, Newman DK. 2004. Phenazines and Other Redox-Active Antibiotics Promote Microbial Mineral Reduction. Applied and Environmental Microbiology 70:921.

35. McRose DL, Newman DK. 2021. Redox-active antibiotics enhance phosphorus bioavailability. Science 371:1033.

36. Yang L, Nilsson M, Gjermansen M, Givskov M, Tolker-Nielsen T. 2009. Pyoverdine and PQS mediated subpopulation interactions involved in Pseudomonas aeruginosa biofilm formation. Molecular Microbiology 74:1380–1392.

37. Braud A, Geoffroy V, Hoegy F, Mislin GLA, Schalk IJ. 2010. Presence of the siderophores pyoverdine and pyochelin in the extracellular medium reduces toxic metal accumulation in Pseudomonas aeruginosa and increases bacterial metal tolerance. Environmental Microbiology Reports 2:419–425.

38. Essar DW, Eberly L, Hadero A, Crawford IP. 1990. Identification and characterization of genes for a second anthranilate synthase in *Pseudomonas aeruginosa*: interchangeability of the two anthranilate synthases and evolutionary implications. J Bacteriol 172:884–900.

39. Stintzi A, Cornelis P, Hohnadel D, Meyer JM, Dean C, Poole K, Kourambas S, Krishnapillai V. 1996. Novel pyoverdine biosynthesis gene(s) of *Pseudomonas aeruginosa* PAO. Microbiology (Reading) 142 ( Pt 5):1181–1190.

40. Huang J, Xu Y, Zhang H, Li Y, Huang X, Ren B, Zhang X. 2009. Temperature-Dependent Expression of phzM and Its Regulatory Genes lasI and ptsP in Rhizosphere Isolate Pseudomonas sp. Strain M18. Applied and Environmental Microbiology 75:6568.

41. Mukherjee S, Moustafa D, Smith CD, Goldberg JB, Bassler BL. 2017. The RhlR quorum-sensing receptor controls Pseudomonas aeruginosa pathogenesis and biofilm development independently of its canonical homoserine lactone autoinducer. PLoS Pathog 13:e1006504.

42. Everett MJ, Davies DT, Leiris S, Sprynski N, Llanos A, Castandet JM, Lozano C, LaRock CN, LaRock DL, Corsica G, Docquier JD, Pallin TD, Cridland A, Blench T, Zalacain M, Lemonnier M. 2023. Chemical Optimization of Selective *Pseudomonas aeruginosa* LasB Elastase Inhibitors and Their Impact on LasB-Mediated Activation in IL-1β in Cellular and Animal infection Models. ACS Infect Dis 9:270–282.

43. Diggle SP, Stacey RE, Dodd C, Cámara M, Williams P, Winzer K. 2006. The galactophilic lectin, LecA, contributes to biofilm development in Pseudomonas aeruginosa. Environ Microbiol 8:1095–104.

44. Chiang P, Burrows LL. 2003. Biofilm formation by hyperpiliated mutants of Pseudomonas aeruginosa. J Bacteriol 185:2374–8.

45. Liberati NT, Urbach JM, Miyata S, Lee DG, Drenkard E, Wu G, Villanueva J, Wei T, Ausubel FM. 2006. An ordered, nonredundant library of Pseudomonas aeruginosa strain PA14 transposon insertion mutants. Proc Natl Acad Sci U S A 103:2833–8.

46. Bisht K, Luecke AR, Wakeman CA. 2022. Temperature-specific adaptations and genetic requirements in a biofilm formed by Pseudomonas aeruginosa. Front Microbiol 13:1032520.

47. Jennings LK, Storek KM, Ledvina HE, Coulon C, Marmont LS, Sadovskaya I, Secor PR, Tseng BS, Scian M, Filloux A, Wozniak DJ, Howell PL, Parsek MR. 2015. Pel is a cationic exopolysaccharide that cross-links extracellular DNA in the Pseudomonas aeruginosa biofilm matrix. Proceedings of the National Academy of Sciences 112:11353.

48. Tetz GV, Artemenko NK, Tetz VV. 2009. Effect of DNase and Antibiotics on Biofilm Characteristics. Antimicrobial Agents and Chemotherapy 53:1204.

49. Brady AJ, Laverty G, Gilpin DF, Kearney P, Tunney M. 2017. Antibiotic susceptibility of planktonic- and biofilm-grown staphylococci isolated from implant-associated infections: should MBEC and nature of biofilm formation replace MIC? Journal of Medical Microbiology 66:461–469.

50. Nielsen CK, Kjems J, Mygind T, Snabe T, Meyer RL. 2016. Effects of Tween 80 on Growth and Biofilm Formation in Laboratory Media. Frontiers in Microbiology 7.

51. Toutain-Kidd CM, Kadivar SC, Bramante CT, Bobin SA, Zegans ME. 2009. Polysorbate 80 Inhibition of Pseudomonas aeruginosa Biofilm Formation and Its Cleavage by the Secreted Lipase LipA. Antimicrobial Agents and Chemotherapy 53:136.

52. Chiang W-C, Pamp SJ, Nilsson M, Givskov M, Tolker-Nielsen T. 2012. The metabolically active subpopulation in Pseudomonas aeruginosa biofilms survives exposure to membrane-targeting antimicrobials via distinct molecular mechanisms. FEMS Immunology & Medical Microbiology 65:245–256.

53. Kaushik KS, Stolhandske J, Shindell O, Smyth HD, Gordon VD. 2016. Tobramycin and bicarbonate synergise to kill planktonic Pseudomonas aeruginosa, but antagonise to promote biofilm survival. npj Biofilms and Microbiomes 2:16006.

54. Herrmann G, Yang L, Wu H, Song Z, Wang H, Høiby N, Ulrich M, Molin S, Riethmüller J, Döring G. 2010. Colistin-Tobramycin Combinations Are Superior to Monotherapy Concerning the Killing of Biofilm Pseudomonas aeruginosa. The Journal of Infectious Diseases 202:1585–1592.

55. Bisht K, Wakeman CA. 2019. Discovery and therapeutic targeting of differentiated biofilm subpopulations. Frontiers in microbiology 10:1908.

56. Hoštacká A, Čižnár I, Štefkovičová M. 2010. Temperature and pH affect the production of bacterial biofilm. Folia Microbiologica 55:75–78.

57. Bremer E, Krämer R. 2019. Responses of Microorganisms to Osmotic Stress. Annual Review of Microbiology 73:313–334.

58. Redelman CV, Maduakolam C, Anderson GG. 2012. Alcohol treatment enhances Staphylococcus aureus biofilm development. FEMS Immunology & Medical Microbiology 66:411–418.

59. Paule A, Roubeix V, Swerhone GDW, Roy J, Lauga B, Duran R, Delmas F, Paul E, Rols JL, Lawrence JR. 2016. Comparative responses of river biofilms at the community level to common organic solvent and herbicide exposure. Environmental Science and Pollution Research 23:4282–4293.

60. Yun HS, Kim Y, Oh S, Jeon WM, Frank JF, Kim SH. 2012. Susceptibility of Listeria monocytogenes Biofilms and Planktonic Cultures to Hydrogen Peroxide in Food Processing Environments. Bioscience, Biotechnology, and Biochemistry 76:2008–2013.

61. Lineback CB, Nkemngong CA, Wu ST, Li X, Teska PJ, Oliver HF. 2018. Hydrogen peroxide and sodium hypochlorite disinfectants are more effective against Staphylococcus aureus and Pseudomonas aeruginosa biofilms than quaternary ammonium compounds. Antimicrobial Resistance & Infection Control 7:154.

62. White-Ziegler CA, Davis TR. 2009. Genome-wide identification of H-NS-controlled, temperature-regulated genes in Escherichia coli K-12. J Bacteriol 191:1106–10.

63. Konkel ME, Tilly K. 2000. Temperature-regulated expression of bacterial virulence genes. Microbes Infect 2:157–66.

64. Hood RD, Peterson SB, Mougous JD. 2017. From Striking Out to Striking Gold: Discovering that Type VI Secretion Targets Bacteria. Cell Host Microbe 21:286–289.

65. Puhar A, Sansonetti PJ. 2014. Type III secretion system. Curr Biol 24:R784–91.

66. Hall S, McDermott C, Anoopkumar-Dukie S, McFarland AJ, Forbes A, Perkins AV, Davey AK, Chess-Williams R, Kiefel MJ, Arora D, Grant GD. 2016. Cellular Effects of Pyocyanin, a Secreted Virulence Factor of Pseudomonas aeruginosa. Toxins 8:236.

67. Lee J, Zhang L. 2015. The hierarchy quorum sensing network in Pseudomonas aeruginosa. Protein Cell 6:26–41.

68. Recinos DA, Sekedat MD, Hernandez A, Cohen TS, Sakhtah H, Prince AS, Price-Whelan A, Dietrich LE. 2012. Redundant phenazine operons in Pseudomonas aeruginosa exhibit environment-dependent expression and differential roles in pathogenicity. Proc Natl Acad Sci U S A 109:19420–5.

69. Stewart PS, Costerton JW. 2001. Antibiotic resistance of bacteria in biofilms. Lancet 358:135–8.

70. Lewis K. 2007. Persister cells, dormancy and infectious disease. Nat Rev Microbiol 5:48–56.

71. Hall CW, Mah TF. 2017. Molecular mechanisms of biofilm-based antibiotic resistance and tolerance in pathogenic bacteria. FEMS Microbiol Rev 41:276–301.

72. Ciofu O, Tolker-Nielsen T. 2019. Tolerance and Resistance of *Pseudomonas aeruginosa* Biofilms to Antimicrobial Agents-How *P. aeruginosa* Can Escape Antibiotics. Front Microbiol 10:913.

73. Çam S, Brinkmeyer R. 2020. The effects of temperature, pH, and iron on biofilm formation by clinical versus environmental strains of Vibrio vulnificus. Folia Microbiologica 65:557–566.

74. Nostro A, Cellini L, Di Giulio M, D’Arrigo M, Marino A, Blanco AR, Favaloro A, Cutroneo G, Bisignano G. 2012. Effect of alkaline pH on staphylococcal biofilm formation. APMIS 120:733–42.

75. Stewart EJ, Ganesan M, Younger JG, Solomon MJ. 2015. Artificial biofilms establish the role of matrix interactions in staphylococcal biofilm assembly and disassembly. Sci Rep 5:13081.

76. Seminara A, Angelini TE, Wilking JN, Vlamakis H, Ebrahim S, Kolter R, Weitz DA, Brenner MP. 2012. Osmotic spreading of Bacillus subtilis biofilms driven by an extracellular matrix. Proceedings of the National Academy of Sciences 109:1116.

77. Rubinstein SM, Kolodkin-Gal I, McLoon A, Chai L, Kolter R, Losick R, Weitz DA. 2012. Osmotic pressure can regulate matrix gene expression in Bacillus subtilis. Molecular Microbiology 86:426–436.

78. D’Souza-Ault MR, Smith LT, Smith GM. 1993. Roles of N-acetylglutaminylglutamine amide and glycine betaine in adaptation of Pseudomonas aeruginosa to osmotic stress. Appl Environ Microbiol 59:473–8.

79. Berry A, DeVault JD, Chakrabarty AM. 1989. High osmolarity is a signal for enhanced algD transcription in mucoid and nonmucoid Pseudomonas aeruginosa strains. J Bacteriol 171:2312–7.

80. Fernandes P, Ferreira BS, Cabral JM. 2003. Solvent tolerance in bacteria: role of efflux pumps and cross-resistance with antibiotics. Int J Antimicrob Agents 22:211–6.

81. Ramos JL, Duque E, Gallegos MT, Godoy P, Ramos-Gonzalez MI, Rojas A, Teran W, Segura A. 2002. Mechanisms of solvent tolerance in gram-negative bacteria. Annu Rev Microbiol 56:743–68.

82. He J, Baldini RL, Deziel E, Saucier M, Zhang Q, Liberati NT, Lee D, Urbach J, Goodman HM, Rahme LG. 2004. The broad host range pathogen Pseudomonas aeruginosa strain PA14 carries two pathogenicity islands harboring plant and animal virulence genes. Proc Natl Acad Sci U S A 101:2530–5.

83. Madsen JS, Lin YC, Squyres GR, Price-Whelan A, de Santiago Torio A, Song A, Cornell WC, Sorensen SJ, Xavier JB, Dietrich LE. 2015. Facultative control of matrix production optimizes competitive fitness in Pseudomonas aeruginosa PA14 biofilm models. Appl Environ Microbiol 81:8414–26.

84. McClure R, Balasubramanian D, Sun Y, Bobrovskyy M, Sumby P, Genco CA, Vanderpool CK, Tjaden B. 2013. Computational analysis of bacterial RNA-Seq data. Nucleic Acids Res 41:e140.

85. Benjamini Y, Hochberg Y. 1995. Controlling the False Discovery Rate: A Practical and Powerful Approach to Multiple Testing. Journal of the Royal Statistical Society: Series B (Methodological) 57:289–300.

86. O’Toole GA. 2011. Microtiter dish biofilm formation assay. J Vis Exp.

87. Zhang L, Fritsch M, Hammond L, Landreville R, Slatculescu C, Colavita A, Mah T-F. 2013. Identification of Genes Involved in Pseudomonas aeruginosa Biofilm-Specific Resistance to Antibiotics. PLOS ONE 8:e61625.

88. Lequette Y, Rollet E, Delangle A, Greenberg EP, Bohin J-P. 2007. Linear osmoregulated periplasmic glucans are encoded by the opgGH locus of Pseudomonas aeruginosa. Microbiology 153:3255–3263.

